# *In silico* analyses on the comparative sensing of SARS-CoV-2 mRNA by intracellular TLRs of human

**DOI:** 10.1101/2020.11.11.377713

**Authors:** Abhigyan Choudhury, Nabarun Chandra Das, Ritwik Patra, Suprabhat Mukherjee

## Abstract

The worldwide outbreak of COVID-19 pandemic caused by SARS-CoV-2 leads to loss of mankind and global economic stability. The continuous spreading of the disease and its pathogenesis takes millions of lives of peoples and the unavailability of appropriate therapeutic strategy makes it much more severe. Toll-like receptors (TLRs) are the crucial mediators and regulators of host immunity. The role of several TLRs in immunomodulation of host by SARS-CoV-2 is recently demonstrated. However, the functionality of human intracellular TLRs including TLR3,7,8 and 9 is still being untested for sensing of viral RNA. This study is hoped to rationalize the comparative binding and sensing of SARS-CoV-2 mRNA towards the intracellular TLRs, considering the solvent-based force-fields operational in the cytosolic aqueous microenvironment that predominantly drive these reactions. Our *in-silico* study on the binding of all mRNAs with the intracellular TLRs shown that the mRNA of NSP10, S2, and E proteins of SARS-CoV-2 are potent enough to bind with TLR3, TLR9, and TLR7 and trigger downstream cascade reactions, and may be used as an option for validation of therapeutic option and immunomodulation against COVID-19.

## 1. Introduction

The worldwide outbreak of Coronavirus disease or COVID-19 pandemic caused by the Severe Acute Respiratory Syndrome Coronavirus-2 (SARS-CoV-2), leads to the infection of about 0.33% of the total world’s population causing the death of 1.24 million people by the first week of November 2020.^1^ The continuous expansion of contagion and pathogenesis, it is taking the shape of another dark age in the history of mankind, not only to health crises but also in bankrupting the global socio-economic status. The SARS-CoV-2 belongs to the β-Coronavirus genus of a 2B group of the Coronaviridae family and is considered the third most virulent type, leading to the highest fatality rate in humans followed by the SARS-CoV and MERS-CoV.^2^ It transmits from person to person mainly via close physical contact and by respiratory aerosols that produces during coughing, sneezing, and even talking within proximity, although some recent studies indicate transmission through fecal matters and fomite-borne contaminations.^3–5^

The proteome of the virus consists of structural proteins, existing in three forms – the Spike ‘S’ protein, the Envelope ‘E’ protein and the Membrane ‘M’ protein. Along with these, 16 forms of non-structural proteins (NSPs) combining together to generate different catalytic models.^6^ The genome, however, is much simpler and comprises of a 29,903 base long, positive-sense single-stranded RNA molecule. This (+)ssRNA genome makes it further feasible to be detectible by the intracellular Toll-like receptors (TLRs) which have a high affinity towards nucleic acid-related pathogen-associated molecular patterns (PAMPs).^7^ Several empirical studies have presented various propositions relating the intracellular TLR 3, 7, 8, and 9 to the cytokine storm produced by the virus that majorly owes it the lethality.^8^ Cytokine storm is apparently the incessant extreme activation of cytokine production leading to prolonged and consistent inflammatory response, which becomes almost continuous due to positive feedback loops in the TLR signaling pathways. The studies are quite successful in relating the role of TLRs in recognizing oligonucleotide PAMPs and triggering the cytokine storm.^9^

The binding of Spike protein with the human ACE2 receptor triggers the pathogenesis of the SARS-CoV-2, leading to the activation of TLRs to activate the proliferation and production of pro-inflammatory cytokines causing cytokine storm, those results in inflammations.^10,11^ From previous studies, it has been found that the spike protein shows binding efficiency with the extracellular domains of TLRs including TLR1, TLR4 and, TLR6, with the strongest affinity with TLR4.^12^ Furthermore, the development of several *in-silico* multi-epitope-based peptide vaccine candidates against the SARS-CoV-2 has shown to be effectively binding with TLR3, TLR4 and, TLR5 to regulate the TLR signaling pathways activation and proliferation.^13^ It has been found that targeting human TLRs, in an order to either block the binding of SARS-CoV-2 or by inhibiting the TLR activation and proliferation that induce the production of pro-inflammatory cytokines using certain TLR agonists, might be used as an effective therapeutic strategy against coronavirus disease.^11^ Additionally, the continuous replication and generation of new virus within the host system indicates that there must be binding efficiency and potency with the intracellular TLRs including TLR3, 7, 8, and 9 in order to induce the severity and pathogenesis of the virus and further introduction to the new host. TLR3 is well known for its sensing capabilities of viral PAMPs and exists as a monomer that is attached to the membrane of endosomes, thereby it detects and binds to certain motifs in the invading viral RNA.^14^ TLR7 expressed majorly in the cerebral cortex, lung, bronchus, breast, kidney, rectum, and smooth muscle tissues, and functions by adhering to the endosome and thereby binding to the viral RNAs with high guanosine and uridine content ^15^ to initiates a MyD88 signal transduction resulting in activation of NF-κB, mitogen-activated protein kinase (MAPK) cascades, as well as IRF-7 and IRF-5 activation via IL-1 receptor-associated kinases (IRAK)-1/2/4 and TNF receptor-associated factor-3/6.^16^ The signaling finally induces the production of pro-inflammatory cytokines including IL-1β, IL-6, IL-12, TNF-α, and IFN-α.^17^ TLR8 is expressed predominantly in the lung and the peripheral blood leucocytes and plays a major role in recognizing GU-rich viral ssRNAs including those of SARS-CoV-2.^18^ TLR9 is predominantly expressed in the spleen, lymph node, bone marrow, and peripheral blood leukocytes and it recognizes CpG motifs in viral DNAs. However, several studies suggest its role in sensing ssRNA fragments generated by the SARS-CoV-2 genome.^19^

However, a lack of precise knowledge regarding the nature of oligonucleotides and their ligating affinity towards the TLRs still pertains to exist. This in context is abstaining the medical research community from developing certain therapeutic interventions that would have been vitally important in this hour of severity. Our study is hoped to rationalize the picture and provide clues regarding the interaction range of the oligonucleotides towards the intracellular TLRs, considering the solvent-based force-fields operational in the cytosolic aqueous microenvironment that predominantly drive these reactions.

## 1. Materials and Methods

### 2.1 Data mining

Current literatures suggest the predominant expression of ten proteins of SARS-CoV-2.^20^ Four of them are structural proteins – the spike protein subunits S1 and S2, the envelope protein (E), and the membrane protein (M). While the other six are non-structural proteins viz. NSP7, NSP8, NSP9, NSP10, papain-like protease (PLpro), and main protease (Mpro). So, first of all, full-length RNA sequences of the aforementioned proteins were retrieved by extensive literature research. Further, the respective sequences were subjected to pairwise alignment with the complete genome of the virus (Accession No. MT438755) available at the GenBank database of NCBI showing ~100% identity matches and 0 gap penalties. From this, it was evident that the quality of the sequences obtained was very high and equivalent to the raw original. Thereafter, crystal structures of TLR3 (PDB ID: 1ZIW) and TLR8 (PDB ID: 5WYX) were obtained from the RCSB PDB database. However, the structures of TLR7 and TLR9 were obtained by performing homology modelling in the SWISS-MODEL server from ExPASy (https://swissmodel.expasy.org/) by using protein sequences obtained from GenBank with Accession No. AAZ99026 and AAZ95518.1 respectively. All the crystal surface structures of TLRs and the whole genome of SARS-CoV-2 were then prepared to visualize (Figure 1).

**Figure 1.**
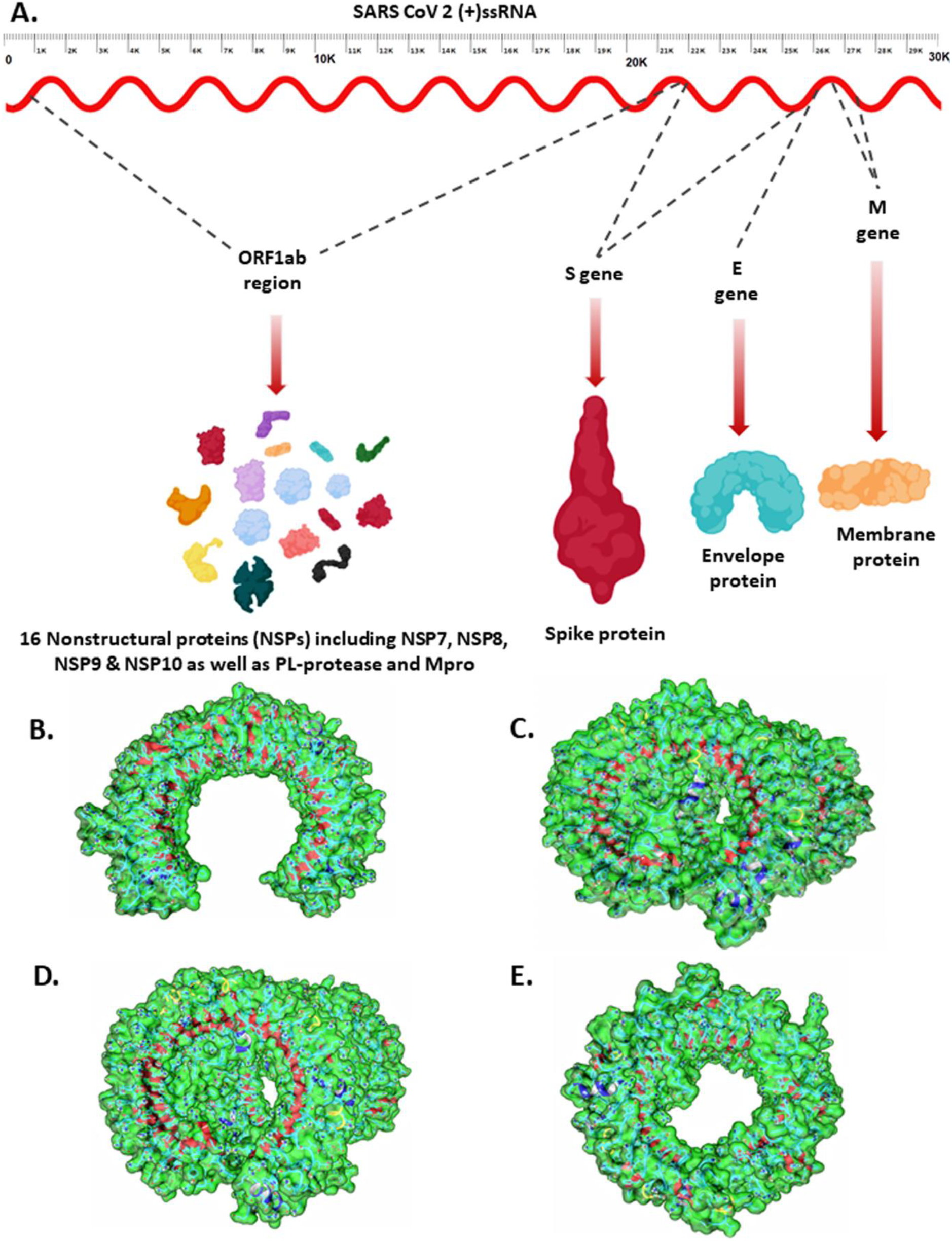
A. Conceptual representation of the SARS-CoV-2 genome. B. TLR3 C. TLR7 D. TLR8 E. TLR9.

To perform further studies, PyMOL is used for removing water molecules, aboriginal heteroatoms as well as any xenobiotic ligands whenever present, and adding polar hydrogens and Kollman charges to the structures.

### 2.2 Prediction of mRNA-Protein interactions

The retrieved RNA sequences were subjected to the imRNA tool developed by IIIT-Delhi (https://webs.iiitd.edu.in/raghava/imrna/). This tool is based on Motif—EmeRging and with Classes—Identification (MERCI) program and scans through the RNA sequences for motifs that are potent to have immunomodulatory properties.^21^ Again, conventional high-throughput experiments for analyzing RNA-protein interactions demand heavier resources and are highly expensive, where RPISeq tool from Iowa State University (http://pridb.gdcb.iastate.edu/RPISeq/) presents a very inexpensive method of predicting RNA-protein interactions, simply by analyzing the RNA and protein sequence data.^22^ RPISeq tool operates on machine learning algorithms and generates outputs in two major forms of RNA-protein interaction prediction parameters – the Random Forest (RF) classifier and Support Vector Machine (SVM) classifier.^23^ Now the protein sequences of TLR3 and TLR8 were extracted from their original PDBs in FASTA format and were used along with the other sequences for the analysis with this tool.

### 2.3 Molecular docking studies and interactions visualization

The HDOCK server (http://hdock.phys.hust.edu.cn/) presents a novel algorithm which is a hybrid of template-dependent along with template-independent *ab initio* free docking. Moreover, it is one of the advance programme which support protein docking against DNA/RNA molecules.^24^ Oligonucleotides have great sizes and demand heavier computing resources for the rendering of molecular models, which is seldomly feasible for any supercomputing server to provide. Thus, HDOCK was a program of choice. The retrieved PDB structures along with the RNA sequences were used for the purpose, however, the species-sensitive and intricate nature of the molecules of interest drove us to go with the template-free docking method. Thereafter, the docked complexes were further visualized with the Protein-Ligand Interaction Profiler (PLIP) tool (https://projects.biotec.tu-dresden.de/plip-web/plip/) it is a Python-based open-source program that provides a complete analysis and visualization of the non-covalent protein-ligand interactions even on single-atom level that include seven prime interaction types *viz*. hydrogen bonds, hydrophobic contacts, π-stacking, π-cation interactions, salt bridges, water bridges, and halogen bonds.^25^

### 2.4 Simulation studies of the docked complexes

#### 2.4.1 Normal mode analysis (NMA) study

The so retrieved docked structures were then fed into the iMOD webserver from ChaconLab (http://imods.chaconlab.org). This program has a user-friendly GUI and is a well-recognized tool for performing normal mode analysis (NMA) and simulating various modal trajectories of protein dynamics. NMA in dihedral coordinates naturally mimics the combined functional motions of protein molecules modelled as a set of atoms connected by harmonic springs.^26^ As output the server delivers affine modelled arrow, vector field and a modal animation to signifies the motions. Moreover, study also provide more detail profile about mobility using B-factor and deformability plots whereas eigenvalue helps to measure the relative modal stiffness of the structure.^27^ Besides, covariance matrix implies the variations among the different mode of motions.

#### 2.4.2 Molecular dynamics (MD) simulation study

In-silico molecular docking between receptor and ligand is not sufficient to conclude the nature of the complex, further analysis viz. MD simulation is desired to validate the structural stability of that complex. GROningen MAchine for Chemical Simulations (GROMACS) is the utmost platform for the MD simulations as it possesses ability to employ different force fields to generate simulation data not only of proteins complexes but also complexes of DNA-protein and RNA-protein.^28^

In this work, Chemistry at Harvard Macromolecular Mechanics (CHARMM) force field was accessed initially to generate necessary input files for MD simulation.^29–31^ Solution builder approach under CHARMM-GUI web tool was first add water box (TIP3 216) and then neutralizing atoms to solvate the system. In order to remove bad contacts and generate more specific outcomes periodic boundary conditions (PBC) were analysed and minimization steps were performed sequentially. Server provided outcomes were then applied on GROMACS v. 2020.1 to achieve the equilibrium (NVT-constant volume and temperature) of the system. Finally, module gmx_mdrun was accessed to run the MD simulation of the system and xmgrace was used to visualize the output of simulation as root mean square deviation (RMSD) plot, radius of gyration (Rg) plot, solvent accessible surface area (SASA) plot and hydrogen bond plot.

### 2.5 Analysis of conformational changes of TLRs during and after MD simulation

Protein with conformational changes from its native form suggests it is in bounded state with any ligand.^32^ Thus, to analyse the changes in TLRs, firstly we extract the PDBs from trajectories and then visualized using PyMOL.

## 3. Results

### 3.1 Homology modelling

In order to assesse the quality of modelled structure of TLR7 and TLR9, we access Structure Assesement tool under SWISS-MODEL server and found out both the structures are significant as more than 92% residues from both structures reside in Ramachandran plot favoured region signifying a stable stereochemical structure (Figure S1).

### 3.2 Motif analysis

Motif analysis played a crucial role in determining the immunoregulatory potency of the given RNA fragments in interacting with TLR proteins and also their role in inciting pro-inflammatory cytokine production.^33^ According to outcomes from imRNA tool Figure S2A was modified where the number of motifs was plotted against the sequence length of the mRNAs to predict the interaction between mRNAs and TLRs. In support RPISeq tool developed predictions on RF and SVM classifier confirm the interactions as they scored above 0.5 and are visualized in Figure S2B-S2F.

### 3.3 Molecular docking studies and interactome analysis

Our study focuses principally on the interactions of the intracellular TLRs with the mRNA fragments, and molecular docking studies play a pivotal role in this study by precisely determining the most probable binding pocket. The docking algorithms used by the HDOCK program globally samples all the binding poses of the RNA to the protein using a fast and flexible Fourier transform (FFT) search strategy with focuses on cavity prediction and shape complementarity.^24^ Further, all the sampled binding modes are analyzed by an iterative knowledge-based scoring function. The more negative the value, the higher the score. Overall, 40 RNA-protein docked complexes were generated in this study and the scores are furnished in Table S1. However, only the four topmost scoring docked complexes were selected for further experimentation and analysis.

#### 3.3.1 Interaction of NSP10 RNA fragment with TLR3

In accord to our docking studies, TLR3 binds with a much proficient binding pose with the RNA fragment encoding NSP10, with a high docking score of −404.77 which is higher than any other docking operation involving TLR3 like that of the RNA fragment of papain-like protease which binds with a score of −346.51 or that of the main protease which has a score of −333.76 (Table S1). While in search of insights of molecular docking PLIP study found the involvement of 251Arg, 252Asn, 277Thr residues in hydrogen bonding, 326Tyr, 359His in π-Cation interactions and Salt bridges from 32 and 319His (Figure 2A, Table 1)

**Table 1.** Residues involved in bonding interactions among the given docked complexes.

| Complex | Interacting residues |  |  |
| --- | --- | --- | --- |
| | Hydrogen bonding residues | π-Cation Interactions | Salt Bridges |
| TLR3 with NSP10 RNA | 251ARG, 252ASN, 277THR, 322PHE | 326TYR, 359HIS | 32HIS, 319HIS |
| TLR7 with E protein RNA | 684LYS, 707SER, 708HIS, 710(B)GLN, 731LYS, 732ASN, 734(B)GLN, 736ARG, 755SER, 756SER, 758LYS, 782HIS, 784(B)ARG | 758(B)LYS, 781HIS, 782(B)HIS | 627ARG, 736ARG, 758(B)LYS, 820(B)HIS |
| TLR8 with NSP8 RNA | 426(B)SER, 470(B)PHE, 473(B)PRO, 489(B)SER, 513(B)SER, 538(B)THR, 562(B)SER, 563TYR, 569ARG, 590(B)ASN, 592(B)SER, 672(B)ASN, 696(B)ARG, 699LYS, 723ARG, 779ASP, 797(B)ARG, 810ARG | 569ARG, 696(B)ARG, 699LYS | 375(B)ARG, 643(B)ARG, 673(B)ASP, 721HIS, 723(B)ARG, 749(B)LYS |
| TLR9 with S2 subunit RNA | 224TYR, 247ARG, 248VAL, 292LYS, 311ARG, 312VAL, 335GLN, 337ARG, 340ASN, 397ARG, 399GLN, 464GLU | 311ARG | 207LYS, 287GLU, 292LYS, 337ARG, 368GLU, 467ARG |

**Figure 2.**
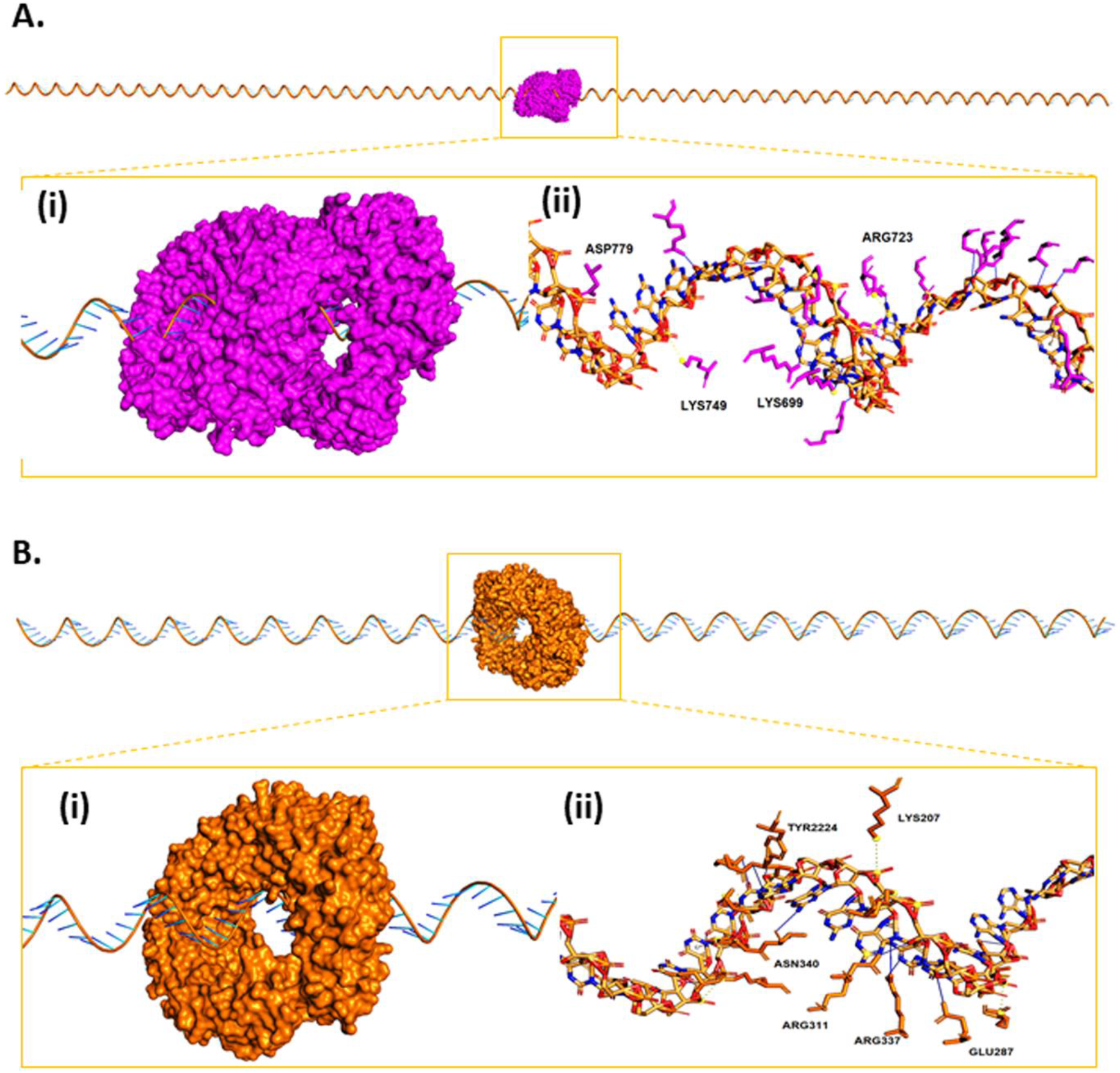
A. Molecular interaction between NSP10 RNA fragment and TLR3 protein. (i) Zoomed in configuration (ii) Detailed bonding interactions determined by PLIP. Solid blue lines represent Hydrogen bonds, dotted orange lines represent π-Cation interactions, while dotted yellow lines denote Salt bridges. B. Molecular interaction between E-protein RNA fragment and TLR7 protein. (i) Zoomed in configuration (ii) Detailed bonding interactions determined by PLIP. Solid blue lines represent Hydrogen bonds, dotted orange lines represent π-Cation interactions, while dotted yellow lines denote Salt bridges. It is to be noted that for the sake of perceptual clarity some residues have been intentionally omitted from the diagram, however the same have been furnished in table 1.

#### 3.3.2 Interaction of Envelope-protein RNA fragment with TLR7

In this study, the docking operations determine a good binding pocket for E-protein mRNA within the TLR7 dimeric cavity, with a binding score of −381.60. PLIP analysis of the docking complex later reveals that the complex stability is regulated by hydrogen bonding interactions through residues *viz.* 684Lys, 707Ser, 708His, by π-Cation Interactions through 758(B)Lys, 781His, 782(B)His residues and through Salt Bridges with residues of 627Arg, 736Arg, 758(B)Lys, 820(B)His as shown in Figure 2B and Table 1.

#### 3.3.3 Interaction of NSP8 RNA fragment with TLR8 protein

Herein, the most probable binding pose of NSP8 RNA fragment and TLR8 is built with a definite score of −416.84, which is much higher than those poses comprising of the RNAs of PL-pro and Mpro which had scores of −360.49 and −355.80 respectively. PLIP analysis as in Figure 3A again confirms these results as it shows that there exists a strong concerted interaction among the complex domains that are 426(B)Ser, 470(B)Phe, 473(B)Pro, 489(B)Ser, 513(B)Ser, etc residues in hydrogen bonding, 569Arg, 696(B)Arg, and 699Lys in π-Cation interactions and Salt bridges from 643(B)Arg, 673(B)Asp, 721His, etc. as shown in Table 1.

**Figure 3.**
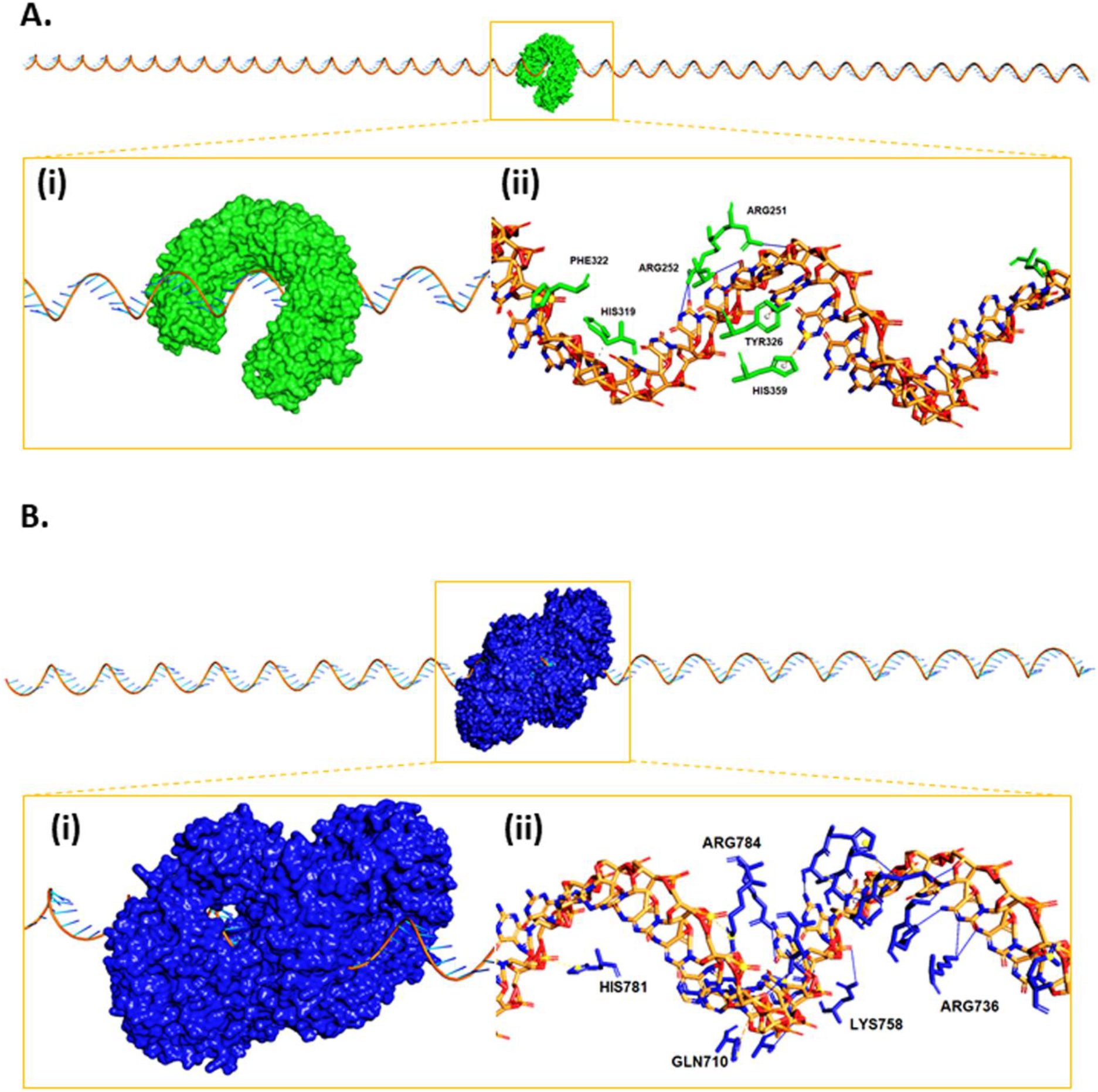
A. Molecular interaction between NSP8 RNA fragment and TLR8 protein. (i) Zoomed in configuration (ii) Detailed bonding interactions determined by PLIP. Solid blue lines represent Hydrogen bonds, dotted orange lines represent π-Cation interactions, while dotted yellow lines denote Salt bridges. B. Molecular interaction between S2-subunit RNA fragment and TLR9 protein. (i) Zoomed in configuration (ii) Detailed bonding interactions determined by PLIP. Solid blue lines represent Hydrogen bonds, dotted orange lines represent π-Cation interactions, while dotted yellow lines denote Salt bridges. It is to be noted that for the sake of perceptual clarity some residues have been intentionally omitted from the diagram, however the same have been furnished in table 1.

#### 3.3.4 Interaction of S2 Subunit RNA fragment with TLR9 protein

Among the docked complexes with TLR9 possessing the S2 subunit mRNA fragment stood at the highest position with a score of −440.33. The molecular insights of this ligation are well supported by the PLIP analysis as finding involves 224Tyr, 247Arg, 248Val, 292Lys residues in hydrogen bonding, 311Arg in π-Cation interactions and Salt bridges from 207Lys, 287Glu, and 292Lys (Figure 3B, Table 1).

### 3.4 Simulation analysis

#### 3.4.1 Normal mode analysis (NMA)

iMODs server generates conformational morphing trajectories of the molecule as depicted in Figure S2G-S2J and enables us to inspect the compatibility and ultimate stability of the complex. Affine-model-based arrow within the structure represents the nature of domain dynamics (Figure S3A, S3B, S4A and S4B). While structure mobility and flexibility are categorized by deformability and B-factor plot (Figure S3C, S3D and Figure S4E, S4F). In deformability plot hinge regions defined the non-rigid helical contents and in B-factor distortions are atomic positional fluctuations. Calculations to measure eigenvalue are fully depended on the energy required to deform the structure, as Figure S4H with very low eigenvalue direct lower energy can deform the structure and infer greater stability. According to Abdelli et al. (2020),^34^ variance shows a direct and inverse relation with eigenvalue as depicted in Figure S3I, S3J, S4I and S4J, where red colour represents individual and green to cumulative variance. In the covariance matrix, motions are categorized in different modes and visualized in Figure S3K and S3M with red, blue and white colour that signify variations among correlated, anti-correlated and uncorrelated motions respectively. In the end, the last outcome of NMA study, elastic network map graph is represented by Figure S4L and S4N where grey dots indicate the stiffness of the motions and springs direct the atomic connections of the complex.

#### 3.4.2 Molecular dynamics (MD) simulation

Four most active complexes, those were primarily strained out according to their significant docking score were subjected to further analysis through MD simulation. MD is the computerized method of simulation that allows mobility of atoms and molecules for a given time to monitor the interaction and calculate the forces, that ultimately figure out the atomic level configuration of the complex structure and then visualize using several graphical plots.^35^ Herein, RMSD plot in Figure 4A reflects the structural stability of the backbone of different receptor TLRs of the four complexes and suggests complex TLR8-NSP8 is less stable among others. Where Figure 4B and 4C define the compactness and solvent accessible area of that complexes as R_g_ plot and SASA plot, respectively. At last the graphical plot of hydrogen bond clearly describes the insights of the complexes and supports the postulation reflected on RMSD, R_g_ and SASA plots (Figure 4D).

**Figure 4.**
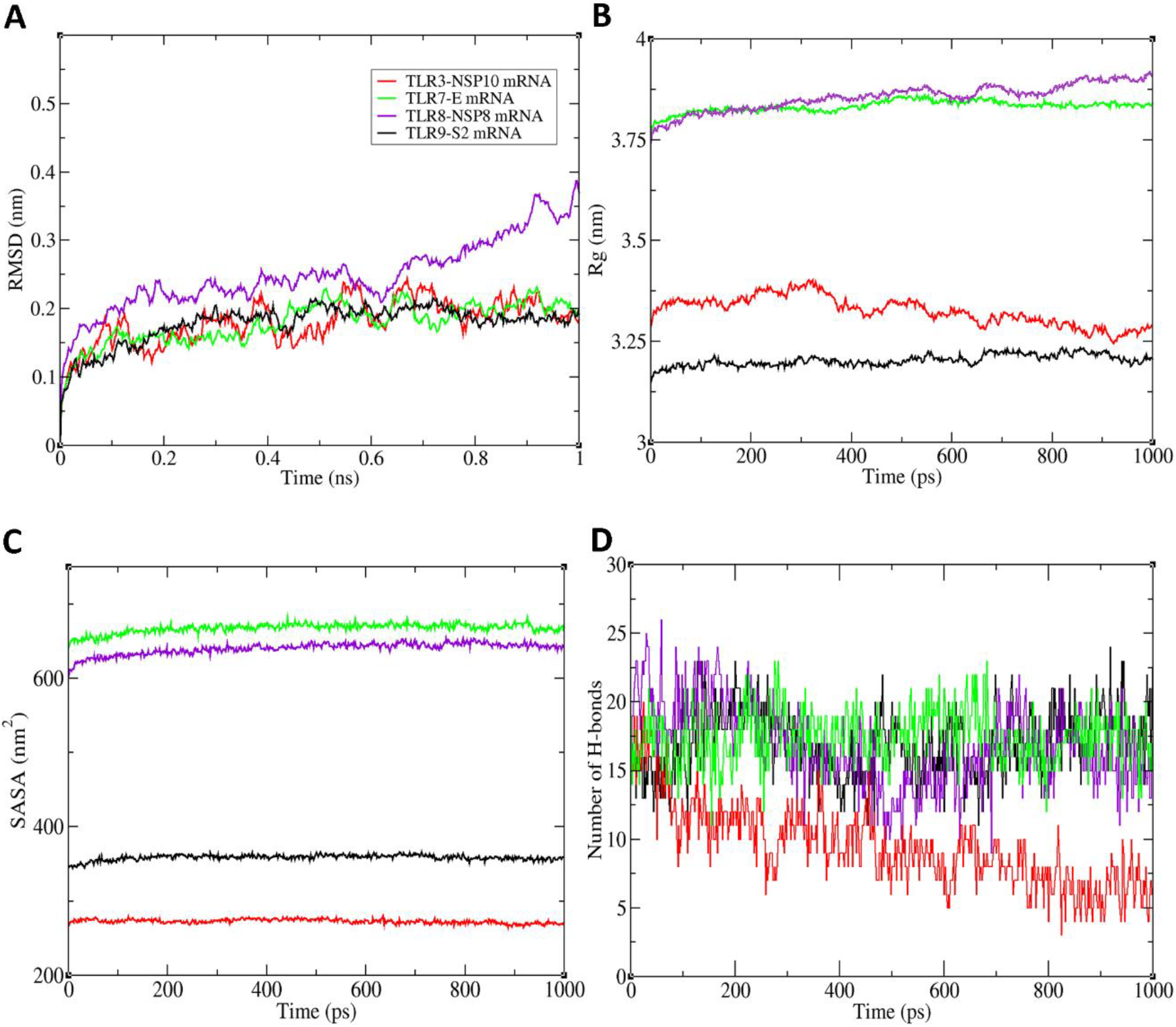
Molecular dynamic (MD) simulation study. Reflecting outcomes generated during MD simulation studies of TLR3-NSP10 mRNA complex, TLR7-E protein mRNA complex, TLR8-NSP8 mRNA complex and TLR9-S2 protein mRNA complex. A. RMSD plots. B. Rg (Radius of gyration) plots. C. SASA plots. D. Graphical presentation of Hydrogen bonds. (TLR3-NSP10 mRNA complex shown in red, TLR8-NSP8 mRNA complex shown in violet, TLR9-S2 protein mRNA complex shown in black and TLR7-E protein mRNA complex in green).

### 3.5 mRNA induced conformational changes in TLRs during and after MD simulation

Apart from RMSD, Rg, SASA and hydrogen bond studies, we also analyse conformational changes of receptor TLRs arisen during simulation processes. Here, we prepared and visualized six different conformation on different time steps of each TLRs and revealed mRNA of NSP10, NSP8, S2 subunit of spike protein and Envelope protein have enough potential to change the conformation of extracellular domain of TLR3, TLR8, TLR9 and TLR7 respectively (Figure 5).

**Figure 5.**
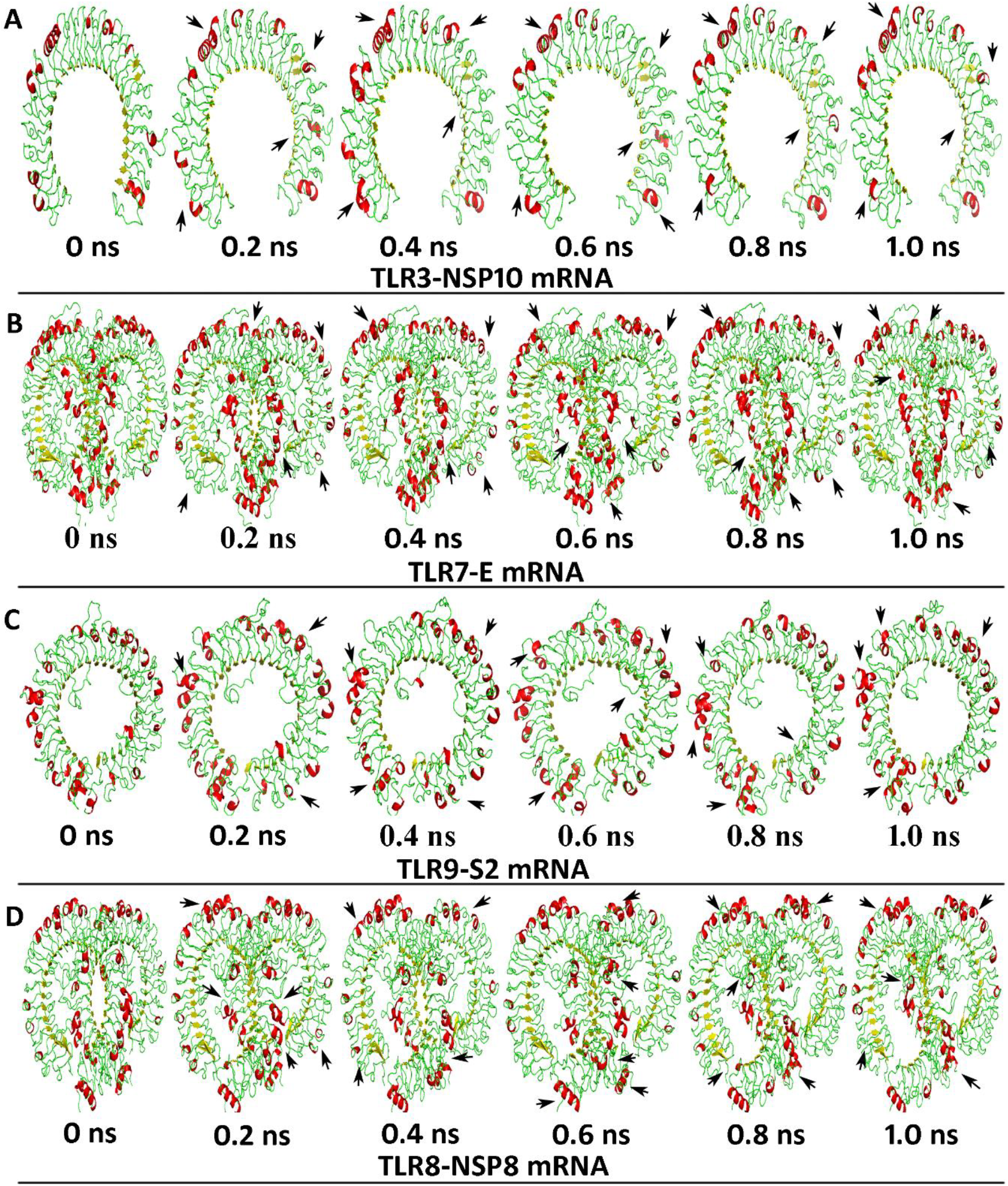
mRNA induced conformational changes in TLR3, TLR7, TLR8 and TLR9. Representing changes in helix, sheet, and loop in the structure of TLRs at various time points of molecular dynamic simulation. A represents changes in TLR3 after NSP10 mRNA induction. B signifies envelope (E) protein mRNA induced alteration of TLR7. C represents changes in TLR9 after S2 subunit of spike protein mRNA induction. While D exhibits NSP8 mRNA induced alteration of TLR8. Red is depicting the helix, yellows are the sheets, and loops are given as green.

## 4. Discussion

Toll-like receptors (TLRs) are the essential mediators and regulators of host immunity. The intracellular TLRs play a vital role in sensing the viral RNA and triggers the downstream activation of signalling cascade of immunomodulation. TLR3 dimerizes upon binding and recruits signal transfer proteins MyD88, TIRAP, TRAM, or TRIF in the cytoplasmic TIR domain.^36^ Certain kinases alike IRAKs, TBK1, and IKKs are activated as well as TRAF6, as per different adaptors, thus leading to activation of the MAPK, NF-κB, or JNK-STAT pathways for promotion of the transcription of inflammatory cytokines and for production of IFN, IL-1β, IL-6, to coordinate the local or systemic inflammatory responses among which IL-1β and IL-6 are the major pro-inflammatory cytokines.^37^ TLR8 protein has various similarities with TLR7 including their gene loci on the X chromosome and believed to possess a close phylogenetic relationship.^38^ The binding to TLR7/8 to the RNA, recruits MyD88 which on one hand activates IRF-7 and on the other hand also activates IRAK-4 resulting in the activation of the transcription factor NF-κB and subsequent antiviral response.^39^ Ligated TLR9 moves from the endoplasmic reticulum to the Golgi apparatus and lysosomes, recruting MyD88-dependent pathway, and resulting in IRF7-mediated IFN production or in proinflammatory responses by producing IL-6, IL-12, and TNFα through activation of the NF-κB.^40^

This study focuses on the establishing the binding and potency of SARS-CoV-2 with the human intracellular TLRs including TLR 3, 7, 8, and 9. Regarding this crystal structures and templates are preliminarily obtained via X-Ray Diffraction techniques with resolution of most 2.50Å, considering good for molecular docking studies. Since the quality of results obtained from further experiments would rely on the stereochemical quality of modelling, and it showed the models had >92% residues in favored regions of their respective Ramachandran plots (Figure S1) indicating standard quality of modelling.

The motif analysis recognizes the immunomodulated motifs within the RNA sequence that can stimulate the immune responses. According to imRNA tool the mRNA fragments are highly capable of interacting with the TLRs and influence immunomodulation (Figure S2A). Though mRNA of the S2 subunit and NSP8 proteins are almost similar in size and have the highest potency compared to others. While RPISeq tool strained out best candidates showing maximum potency in interaction for intracellular TLR 3, 7, 8 and 9. As RPISeq tool predicted a high interaction coefficient for NSP10 RNA with TLR3 as exhibited in Figure S2B. While the threshold SVM and RF classifier values for minimum interaction is 0.5 for SVM classifier as well as for RF classifier, in contrast, the NSP10 mRNA with SVM classifier value of 0.984 and RF classifier value of 0.8 it stands among the best candidates for interacting with TLR3. While prediction figure out E-protein RNA fragment is best in interact TLR7 with an RF classifier value of 0.75 and an SVM classifier value of 0.884 (Figure S2C). Later on, studies also revealed best interaction between NSP8 mRNA and TLR8 protein having SVM and RF classifier as high as 0.991 and 0.7 respectively (Figure S2D). Moreover, prediction again showed a higher probability of interaction of TLR9 with the S2 subunit with SVM classifier value of 0.868 and RF classifier value of 0.7 (Figure S2E).

Further, the molecular docking studies showed a similar trend by reflecting highest docking score to the complexes selected through RPISeq tool and in turn support the prediction of best docked structures (Table S1). After getting confirmation on best predicted structure through HDOCK we forwarded to next step of simulation to analyze the conformational stability and flexibility of that selected four structures viz. TLR8-NSP8 mRNA complex (Figure 3A), TLR3-NSP10 mRNA complex as shown in Figure 2A, TLR7-E mRNA complex in Figure 2B and TLR9-S2 mRNA complex (Figure 3B). Initially, the NMA study reveals all the compounds have better stability as it reflects quite similar and significant eigenvalue scores, those are 3.154899×10^−5^ for TLR8-NSP8 mRNA complex as shown in Figure S4G, 1.27056×10^−5^ TLR3-NSP10 mRNA complex (Figure S3G), 7.042567×10^−5^ TLR7-E mRNA complex as in Figure S3H, and 9.131746×10^−5^ TLR9-S2 mRNA complex (Figure S4H). Later, MD simulation studies figure out postulation of NMA studies is not fully accepted as TLR8-NSP8 mRNA complex found as unstable throughout the simulation process in the RMSD plot Figure 4A. The PLIP analysis Table 1 and hydrogen bond plot Figure 4D finally reveals the interactions between the mRNA ligand and the TLR proteins and the reasons for the stability of the complexes. At the end, our final experiment on changes of conformation of TLRs after binding and simulation hypothesized as mRNA of NSP10, S2 and E proteins are potent enough to bind with TLR3, TLR9 and TLR7 and trigger downstream cascade reactions (Figure 5).

## 5. Conclusion

The intracellular TLRs play a vital role in the recognition of PAMPs and viral pathogens and it is of utmost necessity to understand the sensing of SARS-CoV-2 viral RNA with the human intracellular TLRs including TLR3,7,8 and 9. The current study primarily focuses on the comparative binding of mRNAs of SARS-CoV-2 with the human intracellular TLRs. Later, NMA study found out stability between complexes with significant values for TLR8-NSP8 mRNA complex, TLR3-NSP10 mRNA complex, TLR7-E mRNA complex, and TLR9-S2 mRNA complex. While, upon MD simulation it is confined that the mRNA of NSP8 has lower potency to make complex with TLR8. In turn significant binding potency of NSP10, S2, and E proteins mRNA with TLR3, TLR9, and TLR7 respectively, help us to conclude our study and we infer complexes of TLR3-NSP10 mRNA, TLR7-E mRNA, and TLR9-S2 mRNA can potentially trigger the downstream activation and proliferation of immune responses and can be used in validation for therapeutic action.

## Acknowledgment

We do acknowledge the efforts of all the doctors, health workers, social workers, scientists, and researchers currently working endlessly against coronavirus disease worldwide. We have kept selected studies as reference due to the limitation in space, but we appreciate all the uncited articles dedicated to the research on coronavirus. RP thanks the Department of Higher Education, Govt. of West Bengal for his Merit-cum-means Scholarship.

## Authors’ Contributions

AC, NCD performed all experiments; AC, NCD, and RP write the manuscript; SM design the experiments, analyses the result, edited the manuscript and supervise the study.

## Supplementary Figures

**Figure S1. Validation of homology modelled structures.** A. Ramachandran plot for modelled TLR7 protein, showing 92.11% residues within favored regions. B. Ramachandran plot for modelled TLR9 protein, showing 93.09% residues within favored regions.

**Figure S2. Motif analysis.** A. imRNA Motif analysis graph of different RNAs. B. RPISeq graph for RNAs & TLR3 C. RPISeq graph for RNAs & TLR7 D. RPISeq graph for RNAs & TLR8 E. RPISeq graph for RNAs & TLR9 F. Legend for graphs. G. H. I. & J. Exhibit different conformations of TLR3, −7, −8, & −9 respectively in a given mode, analyzed my iMOD tool.

**Figure S3. Normal mode analysis (NMA) depicting stability and flexibility of complexes as computed by iMODS webserver**. Results of NMA of TLR3-NSP10 mRNA and TLR7-E mRNA complex respectively (A,B) NMA mobility (arrow field), (C,D) deformability, (E,F) B-factor, (G,H) eigenvalue, (I,J) variance (red colour represent individual variances and green colour indicates cumulative variances), (K,M) co-variance map (red-correlate, white-uncorrelated and blue-anti-correlated) and (L,N) elastic network (darker gray regions indicates more stiffer regions) of the complex.

**Figure S4. Normal mode analysis of the TLR-mRNA interactions.** iMODS webserver depicted outcome of NMA of TLR8-NSP8 mRNA and TLR9-S2 mRNA complex respectively, (A,B) NMA mobility (arrow field), (C,D) deformability, (E,F) B-factor, (G,H) eigenvalue, (I,J) variance (red colour represent individual variances and green colour indicates cumulative variances), (K,M) co-variance map (red-correlate, white-uncorrelated and blue-anti-correlated) and (L,N) elastic network (darker gray regions indicates more stiffer regions) of the complex.

## Supplementary table

**Table S1.** Docking scores of different RNA fragments docking with different TLRs.

