## Supplementary material for "*In silico* analyses on the comparative sensing of SARS-CoV-2 mRNA by intracellular TLRs of human"

### **Supplementary Materials**

**Table S1.** Docking scores of different RNA fragments docking with different TLRs.

| **RNA Fragment** | **Docking scores with different TLR proteins** | | | |
|---|---|---|---|---|
| | TLR3 | TLR7 | TLR8 | TLR9 |
| E-protein | -391.41 | -381.60 | -375.33 | -389.97 |
| M-protein | -384.85 | -349.57 | -383.31 | -402.56 |
| Main Protease | -333.76 | -380.90 | -355.80 | -369.86 |
| Papain-like Protease | -346.51 | -342.42 | -360.49 | -367.91 |
| NSP7 | -399.95 | -357.30 | -405.38 | -438.23 |
| NSP8 | -356.90 | -309.00 | -416.84 | -416.66 |
| NSP9 | -378.86 | -329.82 | -358.49 | -431.22 |
| NSP10 | -404.77 | -309.54 | -409.64 | -427.45 |
| S1-subunit | -375.09 | -328.70 | -360.94 | -425.03 |
| S2-subunit | -374.90 | -322.58 | -407.48 | -440.33 |

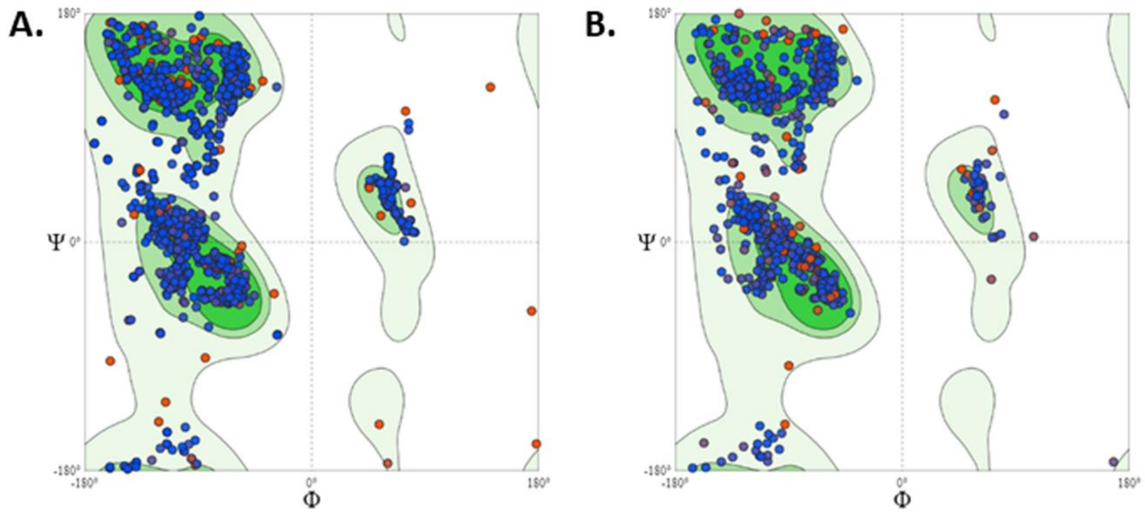


**Figure S1. Validation of homology modelled structures.** A. Ramachandran plot for modelled TLR7 protein, showing 92.11% residues within favored regions. B. Ramachandran plot for modelled TLR9 protein, showing 93.09% residues within favored regions.

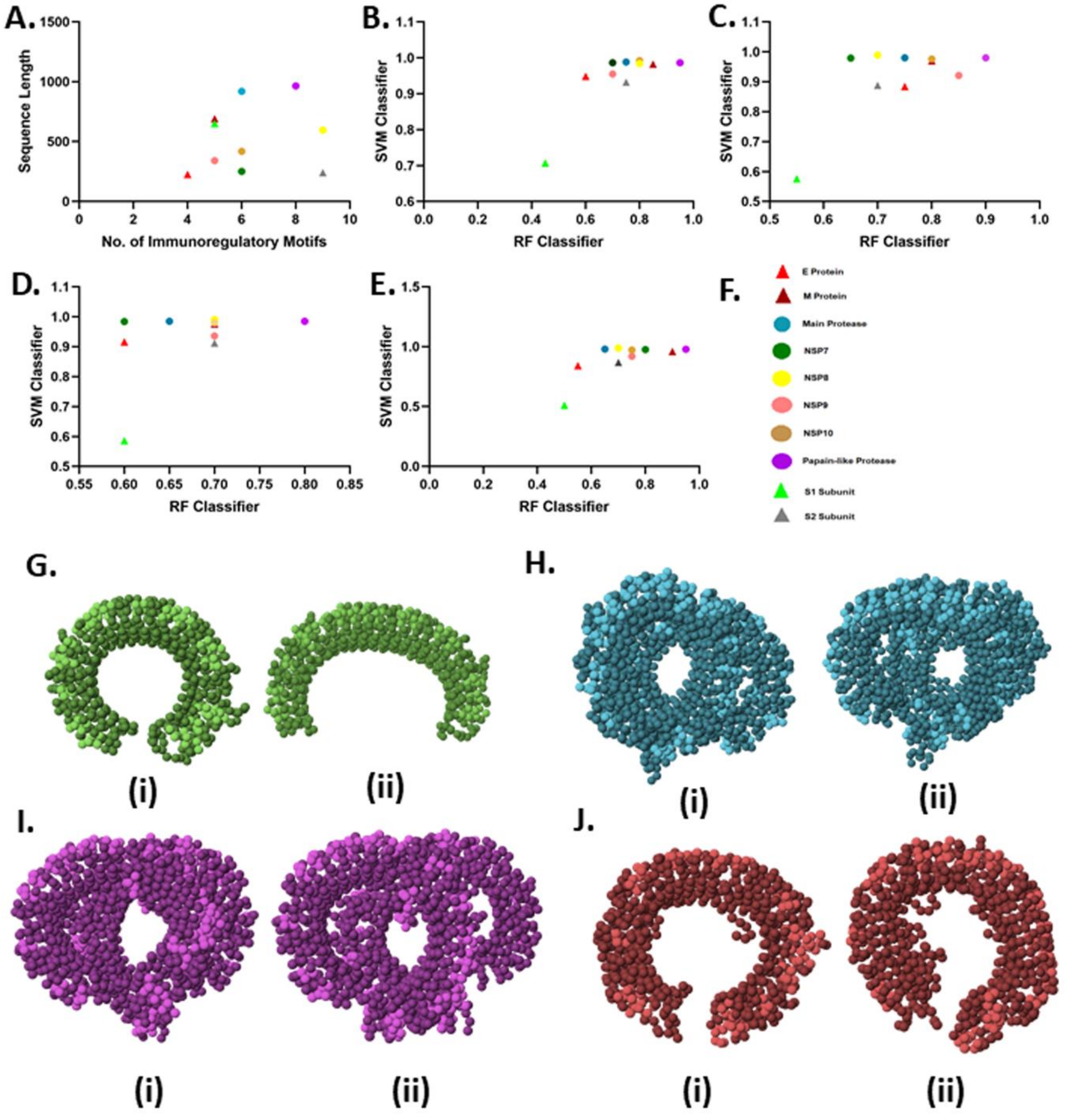


**Figure S2. Motif analysis.** A. imRNA Motif analysis graph of different RNAs. B. RPISeq graph for RNAs & TLR3 C. RPISeq graph for RNAs & TLR7 D. RPISeq graph for RNAs & TLR8 E. RPISeq graph for RNAs & TLR9 F. Legend for graphs. G. H. I. & J. Exhibit different conformations of TLR3, -7, -8, & -9 respectively in a given mode, analyzed my iMOD tool.

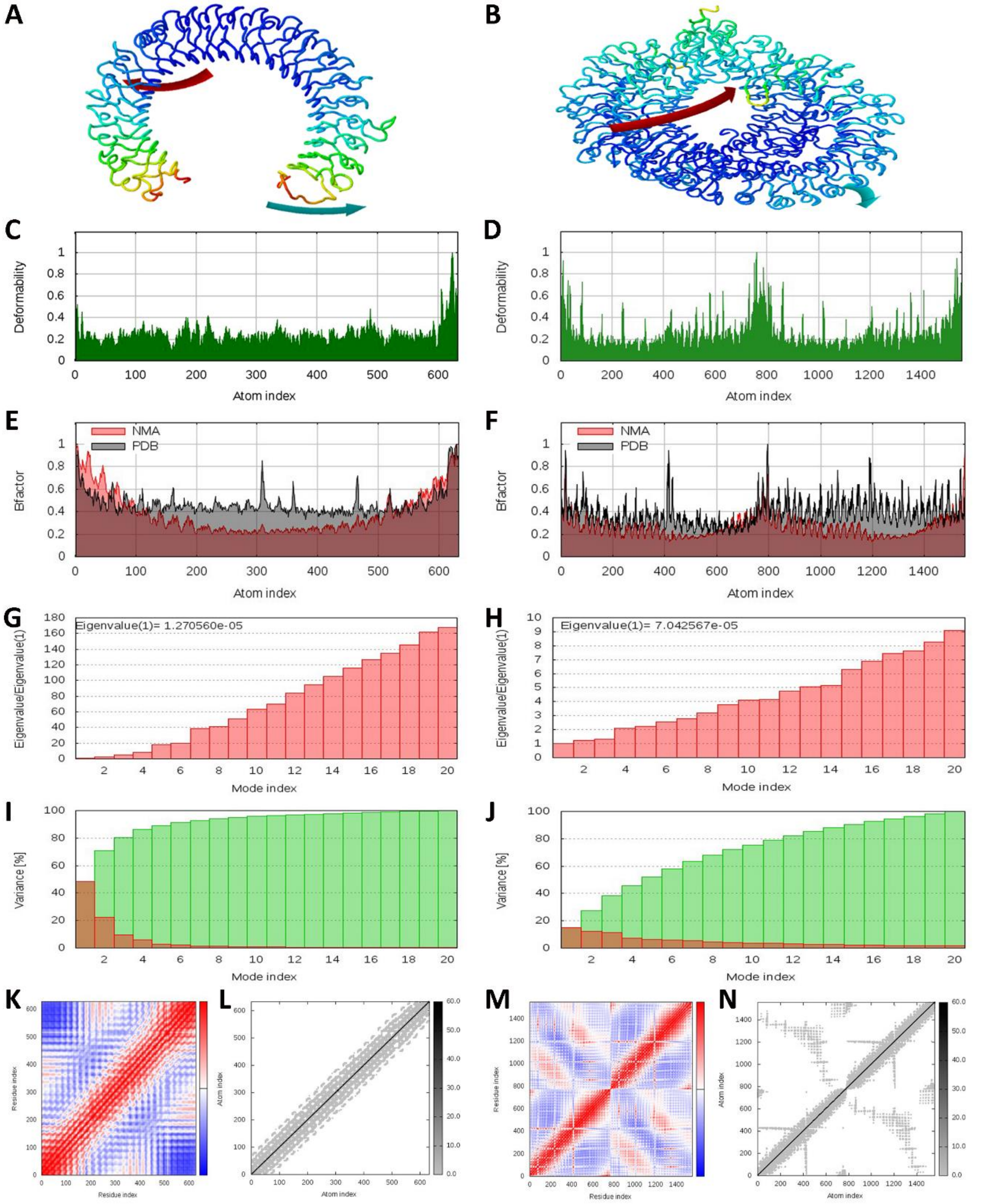


**Figure S3. Normal mode analysis (NMA) depicting stability and flexibility of complexes as computed by iMODS webserver**. Results of NMA of TLR3-NSP10 mRNA and TLR7-E mRNA complex respectively (A,B) NMA mobility (arrow field), (C,D) deformability, (E,F) B-factor, (G,H) eigenvalue, (I,J) variance (red colour represent individual variances and green colour indicates cumulative variances), (K,M) co-variance map (red- correlate, whiteuncorrelated and blue- anti-correlated) and (L,N) elastic network (darker gray regions indicates more stiffer regions) of the complex.

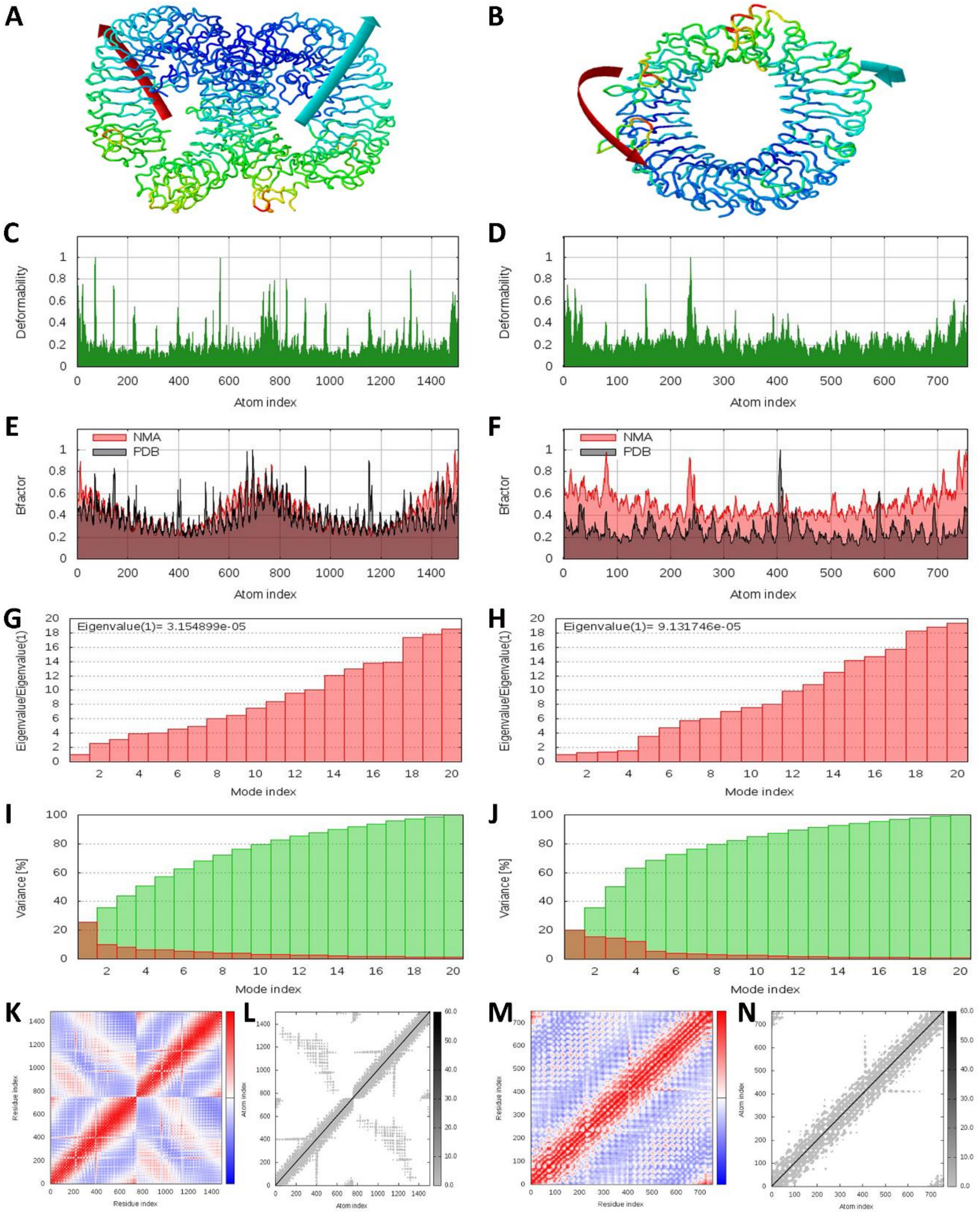


**Figure S4. Normal mode analysis of the TLR-mRNA interactions.** iMODS webserver depicted outcome of NMA of TLR8-NSP8 mRNA and TLR9-S2 mRNA complex respectively, (A,B) NMA mobility (arrow field), (C,D) deformability, (E,F) B-factor, (G,H) eigenvalue, (I,J) variance (red colour represent individual variances and green colour indicates cumulative variances), (K,M) co-variance map (red- correlate, white- uncorrelated and blue- anti-correlated) and (L,N) elastic network (darker gray regions indicates more stiffer regions) of the complex.
